# Loss of endothelial glucocorticoid receptor accelerates diabetic nephropathy

**DOI:** 10.1101/2020.06.29.178228

**Authors:** Swayam Prakash Srivastava, Han Zhou, Ocean Setia, Alan Dardik, Carlos Fernandez-Hernando, Julie Goodwin

## Abstract

Endothelial cells play a key role in the regulation of disease and other developmental processes. Defective regulation of endothelial cell homeostasis may cause mesenchymal activation of other endothelial cells by autocrine effects or of neighboring cell types by paracrine effects, and in both cases contribute to organ fibrosis. However, regulatory control of endothelial cell homeostasis, is not well studied. Diabetes induced renal fibrosis in endothelial GR knock out mice (*GR*^*fl/fl*^;*Tie 1 Cre; GR*^*ECKO*^) but not in control mice (*GR^fl/fl^*); hypercholesterolemia further enhanced severe renal fibrosis in diabetic *GR*^*ECKO*^; *Apoe*^*−/−*^ (DKO) but not in diabetic littermates (*GR*^*fl/fl*^; *Apoe*^*−/−*^). The fibrogenic phenotype in the kidneys of diabetic GR^ECKO^ and diabetic DKO were associated with aberrant cytokine and chemokine reprogramming. Canonical Wnt signaling was identified as new target for the action of endothelial GR. Wnt inhibiton improved kidney fibrosis by mitigating endothelial-to-mesenchymal transition (EndMT) and epithelial-to-mesenchymal transitions (EMT). Similarly, activation of fatty acid oxidation also suppressed kidney fibrosis. Conditioned media from endothelial cells from diabetic GR^ECKO^ stimulated Wnt signaling-dependent epithelial-to-mesenchymal transition in tubular epithelial cells from diabetic controls. These data demonstrate that endothelial GR is an essential antifibrotic core molecule in diabetes.

## Materials and Methods

### Reagents and antibodies

Rabbit polyclonal anti-GR (Cat:SAB4501309) and mouse monoclonal anti-αSMA (Cat:A5228) antibodies were from Sigma (St Louis, MO). Anti-TGFβR1 (Cat:ab31013) antibody was purchased from Abcam (Cambridge, UK). Mouse anti-β-catenin antibody (Cat:610154) was purchased from BD Biosciences. Anti-fibroblast specific protein (FSP1, displayed as S100A4; Cat: 370003) was purchased from Biolegend, CA. Fluorescence-, Alexa Fluor 647-, and rhodamine-conjugated secondary antibodies were obtained from Jackson ImmunoResearch (West Grove, PA). TGFβ2, IL-1β and recombinant TNFα and TGFβ neutralizing antibody were purchased from PeproTech (Rocky Hill, NJ).

### Animal Experimentation

All experiments were performed according to a protocol approved by the Institutional Animal Care and Use Committee at the Yale University School of Medicine and were in accordance with the National Institute of Health (NIH) Guidelines for the Care of Laboratory Animals. Mice lacking the endothelial glucocorticoid receptor (GR) (known as GR^ECKO^) and those lacking this receptor on the Apo E null background (DKO) were generated as previously described^1^. The induction of diabetes in CD-1 mice and C57B/L6 mice was performed according to previously established experimental protocols^2–6^. Briefly, diabetes was induced in 10-week-old GR^ECKO^ mice with five consecutive intraperitoneal (IP) doses of streptozotocin (STZ) 50 mg/kg in 10 mmol/L citrate buffer (pH 4.5). Wnt inhibitor (LGK974) was provided to GR^ECKO^ and control littermate at 5 mg/kg at a frequency of six doses per week for 8 weeks^7^. Etomoxir (20 mg/kg) and c75 (15 mg/kg) were dosed (ip) three times per week for 3 weeks in the GR^ECKO^ and control littermate. A single IP dose of 200 mg/kg STZ was used to induce diabetes in CD-1 mice. Fenofibrate (100 mg/kg), simvastatin (40 mg/kg), were dosed orally for 4 weeks in diabetic CD-1 mice. All mice were sacrificed after 4 weeks of treatment and tissues and blood were harvested. Urine albumin levels were assayed using a Mouse Albumin ELISA Kit (Exocell, Philadelphia, PA).

### Mouse model of unilateral ureteral obstruction (UUO)

UUO surgery procedure was performed as previously described^8^. Briefly, mice were anesthetized with isoflurane (3%–5% for induction and 1%–3% for maintenance). Mice were shaved on the left side of the abdomen, a vertical incision was made through the skin with a scalpel, and the skin was retracted. A second incision was made through the peritoneum to expose the kidney. The left ureter was ligated twice 15 mm below the renal pelvis with surgical silk, and the ureter was then severed between the 2 ligatures. Then, the ligated kidney was placed gently back into its correct anatomical position, and sterile saline was added to replenish loss of fluid. The incisions were sutured and mice were individually caged. Buprenorphine was used as an analgesic. The first dose was administered 30 minutes before surgery and then every 12 h for 72 h, at a dose of 0.05 mg/kg subcutaneously. Mice were sacrificed and kidney and blood samples were harvested after perfusion with PBS at 10 days after UUO. Contralateral kidneys were used as a nonfibrotic control for all experiments using this model.

### Lipid Analysis

Mice were fasted for 12-15 hours and blood was collected by retro-orbital venous puncture. Whole blood was spun down and plasma stored at −80° C. Total cholesterol and triglyceride levels were measured enzymatically by kits from Wako and Sigma, respectively, according to the manufacturer’s instructions.

### Morphological Evaluation

We utilized a point-counting method to evaluate the relative area of the mesangial matrix. We analyzed PAS-stained glomeruli from each mouse using a digital microscope screen grid containing 540 (27× 20) points. Masson’s trichrome-stained images were evaluated by ImageJ software, and the fibrotic areas were estimated.

### Sirius red staining

Deparaffinized sections were incubated with picrosirius red solution for 1 hour at room temperature. The slides were washed twice with acetic acid solution for 30 seconds per wash. The slides were then dehydrated in absolute alcohol three times, cleared in xylene, and mounted with a synthetic resin. Sirius red staining was analyzed using ImageJ software, and fibrotic areas were quantified.

### Immunohistochemistry

Paraffin-embedded kidney sections (5 μm thick) were deparaffinized and rehydrated (2 min in xylene, four times; 1 min in 100% ethanol, twice; 1 min in 95% ethanol; 45 s in 70% ethanol; and 1 min in distilled water), and the antigen was retrieved in a 10 mM citrate buffer pH 6 at 98 °C for 60 min. To block the endogenous peroxidase, all sections were incubated in 0.3% hydrogen peroxide for 10 min. The immunohistochemistry was performed using a Vectastain ABC Kit (Vector Laboratories, Burlingame, CA). Mouse anti-β-catenin antibody (1:100) and CPT1a (Abnova; H00001374-DO1P; 1:100) antibody was used. In the negative controls, the primary antibody was omitted and replaced with the blocking solution.

### Immunofluorescence

Frozen kidney sections (5 μm) were used for immunofluorescence; double positive labeling with CD31/αSMA, CD31/TGFβR1 and E-cadherin/αSMA was measured. Briefly, frozen sections were dried and placed in acetone for 10 min at − 30 °C. Once the sections were dried, they were washed twice in phosphate-buffered saline (PBS) for 5 min and then blocked in 2% bovine serum albumin/PBS for 30 min at room temperature. Thereafter, the sections were incubated in primary antibody (1:100) for 1 h and washed in PBS (5 min) three times. Next, the sections were incubated with the secondary antibodies for 30 min, washed with PBS three times (5 min each), and mounted with mounting medium with DAPI (Vector Laboratories, Burlingame, CA). The immune-labeled sections were analyzed by fluorescence microscopy. For each mouse, original magnification of ×400 pictures were obtained from six different areas, and quantification was performed.

### EndMT and EMT detection

Frozen sections (5 μm) were used for the detection of EndMT and EMT. Cells undergoing EndMT were detected by double-positive labeling for CD31 and αSMA and/or TGFβR1. Cells undergoing EMT were detected by double-positive labeling for E-cadherin and αSMA. Sections were analyzed and quantified by fluorescence microscopy.

### Isolation of endothelial cells

Endothelial cells from the kidneys of non-diabetic and diabetic mice were isolated using a standardized kit (Miltenyl Biotech, USA) by following the manufacturer’s instructions. Briefly, kidneys were isolated and minced into small pieces. Using a series of enzymatic reactions by treating the tissue with trypsin and Collagenase type I solution, a single cell suspension was created. The pellet was dissolved with CD31 magnetic beads and the CD31-labelled cells were separated on a magnetic separator. The cells were further purified on a column. Cell number was counted by hemocytometer and cells were plated on 0.1% gelatin coated Petri dishes.

### Isolation of kidney TECs

After sacrifice kidneys from diabetic GR^ECKO^ and control littermate were excised and perfused with (10 mL) followed by collagenase type II digestion (2 mg/mL). After digestion, the cortical region of kidneys was used for further processing. the cortical region of kidneys was minced and further digested in collagenase buffer for an additional 5 minutes at 37°C with rotation to release cells. Digested tissue and cell suspension were passed through a 70-μm cell strainer, centrifuged at 50 g for 5 min, and washed in PBS for 2 rounds to collect TECs. Isolated TECs were seeded onto collagen-coated Petri dishes and cultured in renal epithelial cell medium (C-26130, PromoCell) supplemented with growth factors for TEC growth.

### Cellular bioenergetic analysis

FAO-associated oxygen consumption rate (OCR) was studied using extracellular flux analysis (Seahorse XFe96, Agilent Technologies). On the assay day, substrate-limited medium was replaced with Krebs-Henseleit buffer assay medium supplemented with 0.2% carnitine for 1h at 37°C without CO_2_. Finally, just before starting the assay, BSA or 200 mM palmitate-BSA FAO substrate was added. After the assay, protein was extracted from wells with 0.1% NP-40–PBS solution and quantified with a bicinchoninic acid protein assay (Thermo Fisher Scientific) for data normalization. OCR was determined as described previously^9^.

### ATP measurement

ATP content was determined using the ATP Colorimetric Assay kit (Biovision), following the manufacturer’s instructions.

### RNA isolation and qPCR

Total RNA was isolated using standard Trizol protocol. RNA was reverse transcribed using the iScript cDNA Synthesis kit (Bio-Rad) and qPCR was performed on a Bio-Rad C1000 Touch thermal cycler using the resultant cDNA, as well as qPCR Master mix and gene specific primers. The list of mouse primers used is given in Table S1.

Results were quantified using the delta–delta-cycle threshold (Ct) method(ΔΔCt). All experiments were performed in triplicate and 18S was utilized as an internal control.

### Western blot

Protein lysates were boiled in sodium dodecyl sulfate (SDS) sample buffer at 94°C for 5 min. After centrifugation at 17,000×*g* for 10 min at 4°C, the supernatant was separated on 6 %-12 % SDS polyacrylamide gels, and blotted onto PVDF membranes (Immobilon, Bedford, MA) via the semidry method. After blocking with TBS (Tris buffered saline containing 0.05 % Tween 20) containing 5 % bovine serum albumin (BSA), membranes were incubated with each primary antibody (GR: 1:1000; Anti-TGFβR1: 1:500; anti-αSMA: 1:500; anti-β-catenin:1:500 and anti-FSP-1: 1:100), in TBS containing 5 % BSA at 4°C overnight. Protein bands were visualized using the Odyssey Infrared Imaging System (LI-COR Biotechnology), and densitometry was performed using ImageJ software (NIH).

### *In vitro* experiments and siRNA transfection

HUVECs were used at passage 4–8 and cultured in Endothelial Basal Medium-2 media with growth factors and 10% serum. Human GR-specific siRNA (Invitrogen) was used at a concentration of 100 nM for 48 h to effectively knock down GR. Cells were treated with or without TGFβ2 (10 ng/ml) for 48 h and harvested for western blot analysis. Some transfected cells were treated with fenofibrate (1μM) and etomoxir (40 μM) for 48 h. In a second set of experiments Human HK-2 cells were cultured in DMEM and Keratinocyte-SFM (1X) medium (Life Technologies Green Island NY). When the cells reached 70% confluence, conditioned media from control siRNA and GR siRNA-transfected HUVECs was added to the HK-2 cell culture.

### Fatty acid uptake

Cultured isolated kidney endothelial cells were incubated with medium containing 2 μCi [^14^C]-palmitate. [^14^C]-palmitate uptake was measured by liquid scintillation counting.

### Fatty Acid Oxidation

Cultured isolated kidney endothelial cells were incubated with medium containing 0.75 mmol/L palmitate (conjugated to 2% free fatty acid–free BSA/[^14^C] palmitate at 2 μCi/mL) for 2 h. One mL of the culture medium was transferred to a sealable tube, the cap of which housed a Whatman filter paper disc. ^14^CO_2_ trapped in the media was then released by acidification of media using 60% perchloric acid. Radioactivity that had become adsorbed onto the filter discs was then quantified by liquid scintillation counting.

### Statistical analysis

All values are expressed as means ± SEM and analyzed using the statistical package for the GraphPad Prism 7 (GraphPad Software, Inc., La Jolla, CA). One-way Anova, followed by Tukey’s test was employed to analyze the significance when comparing multiple independent groups. The post hoc tests were run only if F achieved P< 0.05 and there was no significant variance in homogeneity. In each experiment, N represents the number of separate experiments (in vitro) and the number of mice (in vivo).

Technical replicates were used to ensure the reliability of single values. Data analysis were blinded. The data were considered statistically significant at P< 0.05.

## Introduction

Approximately one-third of diabetic patients will develop diabetic nephropathy (DN), a leading cause of end-stage renal disease (ESRD) that causes more than 950,000 deaths globally each year^10,11^. Over the last two decades, no new drugs have been approved to specifically prevent diabetic nephropathy or to improve kidney function^12^. This lack of advancement stems, in part, from poor understanding of the mechanism of progressive kidney dysfunction. Thus, this knowledge gap contributes to suboptimal treatment options available for these patients. Improved understanding of mechanisms of pathogenesis of diabetic kidney disease is urgently needed to catalyze the development of novel, effective and safe therapeutics which can be targeted to the early stages of diabetes, before kidney damage becomes irreversible.

Diabetic nephropathy is characterized by excess deposition of extracellular matrix, loss of capillary networks and accumulation of fibrillary collagens, activated myofibroblasts and inflammatory cells^13^. In renal fibrosis, myofibroblasts are believed to be an activated fibroblast phenotype that contributes to fibrosis^14^. There are six well-reported sources of matrix-producing myofibroblasts: (1) activated residential fibroblasts, (2) differentiated pericytes, (3) recruited circulating fibrocytes, (4) those from macrophages via macrophage-to-mesenchymal transition (5) those from mesenchymal cells derived from tubular epithelial cells via epithelial-to-mesenchymal transition (EMT) and (6) those from mesenchymal cells transformed from endothelial cells (ECs) via endothelial-to-mesenchymal transition (EndMT)^15,16^,.

Among these diverse origins of matrix-producing fibroblasts, mesenchymal cells transformed from EC via EndMT^15,16^, are an important source of myofibroblasts in several organs, including the kidney^17^. EndMT is characterized by the loss of endothelial markers, including cluster of differentiation 31 (CD31), and acquisition of the expression of mesenchymal proteins including α-smooth muscle actin (αSMA), vimentin, and fibronectin proteins^15,16^.

EC are critical contributors to the formation of new blood vessels in health and life-threatening diseases^18^. Disruption in the central metabolism of EC contributes to disease phenotypes^19,20^. Carnitine palmitoyltransferase 1a (CPT1a)-mediated fatty acid oxidation (FAO) regulates the proliferation of EC in the stalk of sprouting vessels^21–23^. EC use metabolites/precursors for epigenetic regulation of their sub-type differentiation and maintain crosstalk through metabolites release with other cell types^18,23^. Notably, EndMT causes alteration of endothelial cell metabolism, and is an area of active investigation^24,25^. For example, mesenchymal cells derived from EndMT reprogram their metabolism and show defective fatty acid metabolism^25^.

The contribution of EndMT to renal fibrosis has been analyzed in several mouse models of chronic kidney disease^8,14,15,26^. Zeisberg et al., performed seminal experiments and reported that approximately 30~50% of fibroblasts co-expressed the EC marker CD31 along with markers of fibroblasts and myofibroblasts such as fibroblast specific protein-1 (FSP-1) and αSMA in the kidneys of mice subjected to unilateral ureteral obstruction nephropathy (UUO)^8^. The complete conversion from EC into mesenchymal cell types is not needed as intermediate cell types are sufficient to cause activation of profibrogenic pathways. EndMT can induce profibrogenic signaling by its autocrine and/or by paracrine manner to neighboring cells thereby contributing to global fibrosis in kidneys^14,27^. Thus, targeting EndMT might have therapeutic potential for the treatment of renal fibrosis^8,14,28^.

The glucocorticoid receptor (GR) is a nuclear hormone receptor that is expressed ubiquitously in most cell types and is important in many states of health and disease. Glucocorticoid receptors (GRs) mediate the action of steroid hormones in a variety of tissues, including the kidney. Our previous work has demonstrated that tissue-specific loss of this receptor can produce profound phenotypes^1,29–31^. The role of glucocorticoids in cardiovascular and kidney disease is complex. Earlier, we identified endothelial GR as a negative regulator of vascular inflammation in models of sepsis^29^ and atherosclerosis^1^. Moreover, we have also demonstrated loss of endothelial GR results in upregulation of the canonical Wnt signaling pathway (Zhou et al, JCI Insight, In press). Notably, this pathway is known to be up regulated in renal fibrosis^32^. However, whether endothelial GR contributes to the regulation of fibrogenic processes during kidney fibrosis is not known. To begin to understand how endothelial GR may be regulating renal fibrosis we studied endothelial specific GR-knock out mice in both diabetic and non-diabetic conditions.

## Results

### Endothelial GR deficiency results in a fibrogenic phenotype in the kidneys of diabetic mice

The streptozotocin (STZ)-induced diabetic CD-1 mouse is the established mouse model to study diabetic kidney disease^2,3,33^, as the kidney fibrosis phenotype is dependent upon mouse strain specificity^33^. Though STZ-induced diabetic CD-1 mice and diabetic C57B/L6 mice demonstrate similar blood glucose levels, the kidneys of diabetic CD-1 mice have been shown to have higher rates of EndMT and more severe fibrosis when compared to the kidneys of diabetic C57B/L6 mice^2,34^. Therefore, diabetic CD-1 mice are considered pro-fibrotic strain while diabetic C57B/L6 mice are considered to be a less-fibrotic strain^34,35^.

CD31-positive cells from diabetic CD-1 mouse kidneys displayed significant suppression of GR compared to those from diabetic C57B/L6 mice as assessed by immunofluorescent staining. **(Fig. 1a)**. Moreover, EC isolated from the kidneys of diabetic CD-1 mice showed dramatic suppression in both the GR protein level and GR mRNA level when compared to the diabetic C57B/L6 mice and the non-diabetic controls of both genotypes **(Fig. 1b)**.

**Figure 1.**
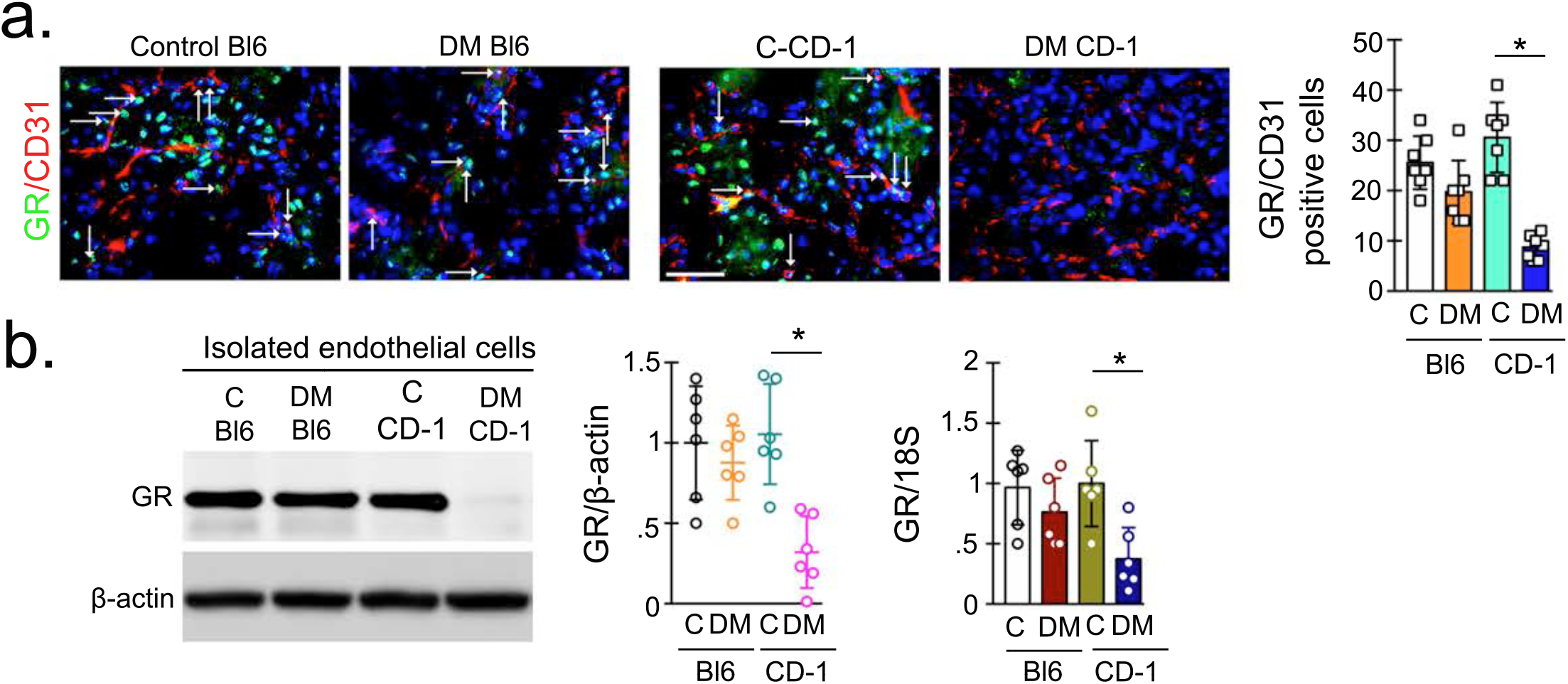
Loss of endothelial GR results in a fibrogenic phenotype in the kidneys of diabetic mice. **(a)** Immunofluorescence analysis was performed in the kidneys of control and diabetic CD-1 and C57Bl6 mice fluorescence microscopy using FITC-labeled GR, Rhodamine-labeled CD31 and DAPI (blue). Merged images are shown. Scale bar: 50 μm in each panel. Representative pictures are shown. N=6/group. Data in the graph are shown as mean ± SEM. **(b)** Western blot and qPCR analysis of GR protein and mRNA levels in isolated endothelial cells from the kidneys of control and diabetic CD-1 mice. Densitometry analysis is normalized to β-actin. mRNA expression is normalized to 18S. N=6/group. Data in the graphs are shown as mean ± SEM. Tukey test was used for analysis of statistical significance. *p <0.05

### Loss of EC GR worsens kidney fibrosis

To verify efficient GR excision from endothelial cells in the kidneys of GR^ECKO^ mice, we performed Western blot and qPCR for GR. As shown in **Fig S1a-b**, mRNA and protein levels were significantly diminished, as expected. Diabetes was induced by injecting 5 consecutive low doses of STZ (50 mg/kg/day IP) in 8-week old GRfl/fl; Tie1 Cre+ (GR^ECKO^) and Cre-littermate controls (GR^fl/fl^) and GR^fl/fl^;Tie1 Cre+/*Apoe*^*−/−*^ (DKO) mice and Cre-littermates (GR^fl/fl^; *Apoe^−/−^*) (**Fig 2a**). Animals were monitored for 4 months post-STZ treatment before sacrifice. At the time of sacrifice, diabetic GR^ECKO^ and diabetic DKO mice and their diabetic littermate controls had no significant change in body weight, blood glucose, heart weight, liver weight, triglycerides or cholesterol; however, diabetic GR^EC KO^ and diabetic DKO had relatively higher kidney weight, spleen weight and albumin-to-creatinine ratios when compared to their respective diabetic controls. **(Fig. 2b-i)**. Diabetic DKO had significantly higher kidney weight and albumin-to-creatinine ratios when compared to diabetic GR^EC KO^. Renal fibrosis was assessed by histologic analysis of kidney sections from all genotypes. Diabetic GR^ECKO^ mice exhibited a higher relative area of fibrosis, higher relative collagen deposition and more severe glomerulosclerosis at the 4-month timepoint when compared to diabetic littermate controls. Diabetic DKO exhibited greatly increased relative area of fibrosis and relative collagen deposition when compared to diabetic ApoE^−/−^ controls and diabetic GR^ECKO^ **(Fig. 2j)**. Immunofluorescence data showed higher collagen I and fibronectin deposition in the kidneys of diabetic animals with GR^ECKO^ with the highest deposition observed in DKO mice **(Fig. 2k-l)**.

**Figure 2.**
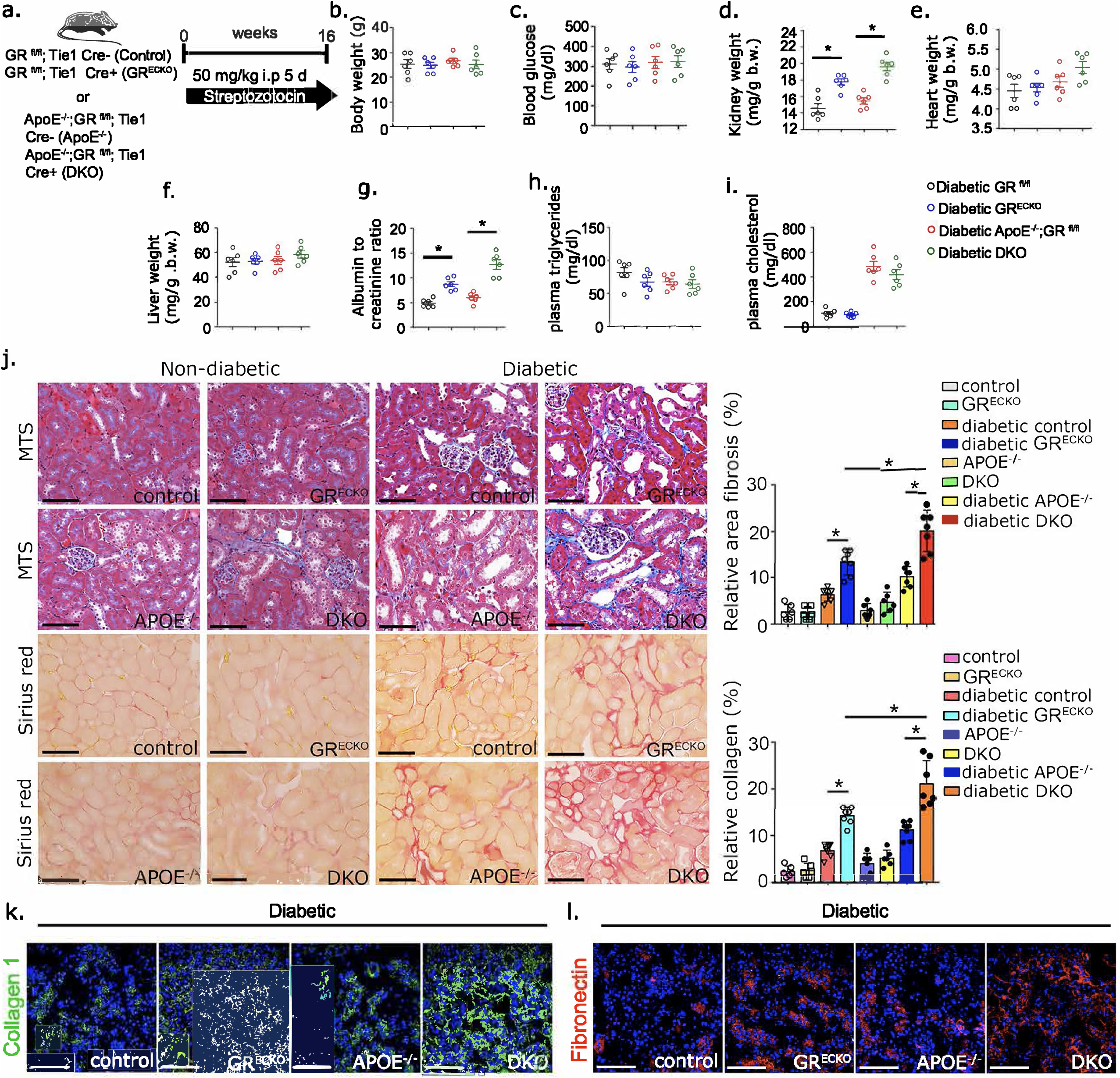
Loss of endothelial GR worsens fibrosis in kidneys of diabetic mice. **(a)** Schematic diagram, showing induction of diabetes in the *GR ^fl/fl^; Tie1 Cre-* (Control) *GR ^fl/fl^; Tie1 Cre+* (GR^ECKO^), *Apoe^−/−^;GR ^fl/fl^; Tie1 Cre-* (*Apoe^−/−^*) and *Apoe^−/−^;GR ^fl/fl^; Tie1 Cre+* (DKO) mice. Five doses of STZ (50 mg/kg/day i.p.) were injected to induce fibrosis. **(b-h)** Physiological parameters including body weight, blood glucose, kidney weight, heart weight, spleen weight, liver weight and albumin-to-creatinine ratio (ACR) were measured. N= 6/group. **(i).** Masson trichome and Sirius red staining in kidneys of non-diabetic and diabetic control, GR^ECKO^, Apoe^−/−^ and DKO were analyzed. Representative images are shown. Relative area fibrosis (%) and relative collagen (%) were measured using the ImageJ program. N=6 non-diabetic group, N=7 diabetic groups. Scale bar: 50 μm in each panel. Data are shown as mean ± SEM. **(j)** Immunofluorescence analysis of collagen I was analyzed in the kidneys of diabetic control, diabetic GR^ECKO^, diabetic ApoE^−/−^, and diabetic DKO with FITC-labeled Collagen 1 and DAPI (blue). Representative images are shown. **(k)** Rhodamine-labeled fibronectin and DAPI (blue). Representative images are shown. Scale bar: 50 μm in each panel. N=6/group. Tukey test was used for the analysis of statistical significance. * p < 0.05.

In order to test the role of endothelial GR in non-diabetic fibrosis, we performed unilateral ureteral obstruction (UUO) in 8-week-old GR^ECKO^ and control littermates **(Fig. S2a)**. There was no significant difference in renal fibrosis between contralateral kidneys of controls and GR^ECKO^ mice. However, UUO kidneys from GR^ECKO^ mice showed a greater relative area fibrosis and greater collagen deposition when compared to UUO kidneys of littermate controls **(Fig. S2b)**. Immunofluorescence staining revealed higher collagen I, αSMA, and fibronectin deposition in the UUO kidneys of GR^ECKO^ when compared to UUO kidneys of control littermates **(Fig. S2c)**.

### Endothelial GR loss reprograms cytokine and chemokine homeostasis

Inflammation is a key factor during the fibroblast activation process in the kidneys of diabetic mice^36,37^ and disruption of cytokine and chemokine homeostasis contributes to the development of diabetic kidney disease^38–40^. To investigate whether there where derangements in homeostasis in our model, we performed cytokine analysis in the plasma of diabetic mice with severe fibrosis (diabetic CD-1) and the plasma of diabetic mice with less severe fibrosis (diabetic C57B/L6). Diabetic CD-1 mice demonstrated higher levels of plasma IL-1β, IL-6, IL-10, IL-17, G-CSF, IFN-γ, TNF-α, MCP-1, CCL3 and CCL4 levels, however the level of CCL5 were remarkably suppressed when compared to that of diabetic C57B/L6 mice **(Fig 3a)**. The same cytokines were also analyzed in the plasma from diabetic GR^ECKO^ mice and littermate controls and diabetic DKO and diabetic *Apoe*^*−/−*^ controls. A similar pattern was observed in both genotypes in that IL-1β, IL-6, IL-10, Eotaxin, G-CSF and CCL4 were significantly higher, while CCL5 was significantly lower, in the plasma of GR^ECKO^ and DKO mice when compared to the plasma of their respective diabetic control littermates **(Fig 3b)**. Similarly, during mRNA gene expression analysis, the level of IL-1β, IL-6, IL-10, IL-17, Eotaxin, and CCL4 were significantly upregulated whereas, CCL5 was significantly downregulated, in the kidneys of diabetic GR^ECKO^ and diabetic DKO mice when compared to the diabetic kidneys of their respective control littermates **(Fig 3c)**, indicating more EC inflammation in mice lacking endothelial GR.

**Figure 3.**
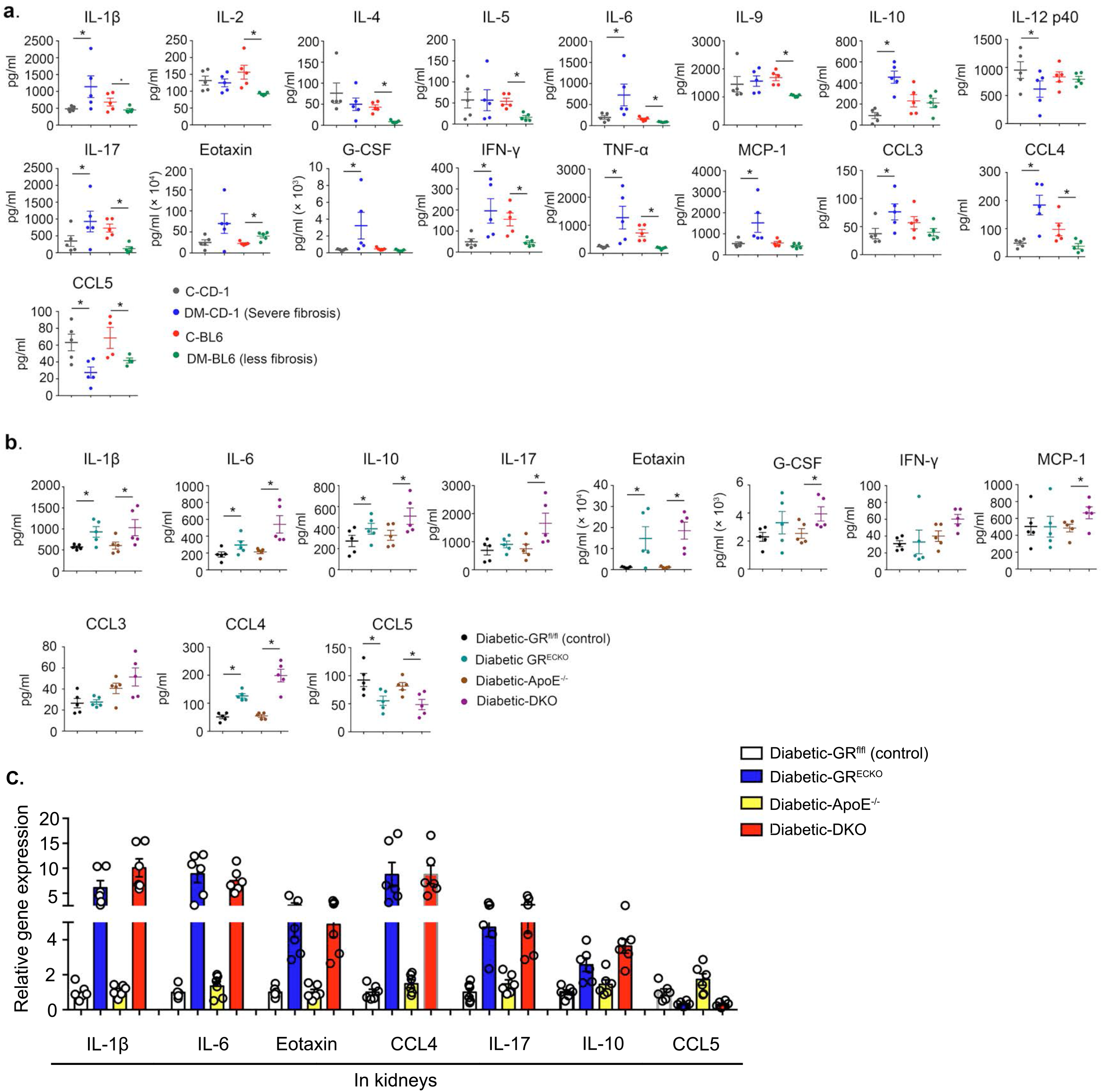
Diabetic kidney disease is associated with cytokine and chemokine reprogramming. **(a)** Cytokines and chemokines were measured in the plasma by using a cytokine array analysis (Luminex). The plasma of control and diabetic mice was analyzed. N=5/group. **(b)** The plasma of non-diabetic and diabetic GR^ECKO^ and DKO mice were analyzed. N=5/group. Data are shown as mean ± SEM. **(c)** Relative gene expression analysis of the indicated molecules in the diabetic kidneys. N=6/group. 18S was used to normalize the expression level. Tukey test was used for the analysis of statistical significance. *p < 0.05.

### Canonical Wnt signaling is a new drug target for the action of endothelial GR

Given the recently described regulation of Wnt signaling by endothelial GR (Zhou et al., JCI Insight, In press), we assessed the mRNA expression of Wnt-dependent genes and fibrogenic markers in EC isolated from the kidneys of diabetic GR^ECKO^ and diabetic DKO mice and their diabetic littermate controls **(Fig. 4a)**. The expression level of Wnt-dependent genes and fibrogenic markers was upregulated in kidneys of diabetic GR^ECKO^ and diabetic DKO when compared to their respective controls. However, the kidneys of diabetic DKO mice showed the highest expression of both Wnt-dependent genes, such as *axin2* and *tcf*, and fibrogenic markers, such as *αSMA* and *fibronectin* as well as the most severe suppression in CD31, suggestive of EndMT. These results were also confirmed at the protein level by Western blotting (**Figure 4b**). Using immunofluorescent co-staining, the same pattern was also observed, with diabetic GR^ECKO^ and diabetic DKO mice demonstrating higher levels of αSMA/CD31 and TGFβR1/CD31 co-staining in the kidneys when compared to their respective controls **(Fig. 4c)**.

**Figure 4.**
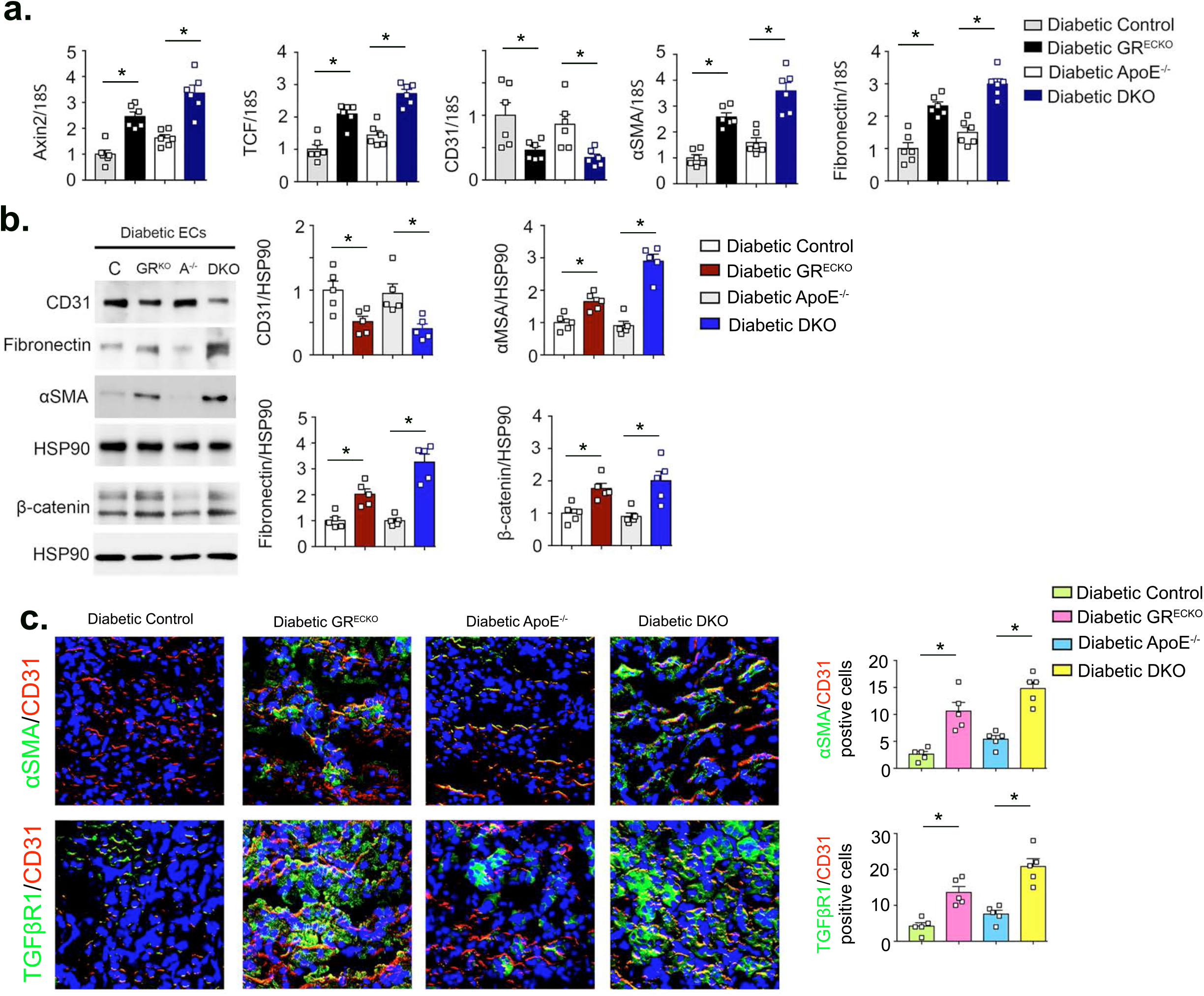
Up regulation of Wnt signaling and fibrogenic markers with loss of endothelial GR. **(a)** Relative mRNA levels determined by qRT-PCR of *Axin2*, *Tcf*, *αSMA*, *CD31* and *fibronectin* were analyzed in isolated EC from the kidneys of diabetic control, GR^ECKO^, *Apoe*^*−/−*^ and DKO mice. N=6/group. **(b)** Western blot analysis of CD31, αSMA, fibronectin, and β-catenin in isolated endothelial cells from the kidneys of diabetic control, GR^ECKO^, *Apoe*^*−/−*^ and DKO mice. N=5/group. Representative blots are shown. Densitometry normalization was performed to HSP90. **(c)** Immuno-fluorescence analysis of αSMA/CD31 and TGFβR1/CD31 was performed in the kidneys of diabetic control, GR^ECKO^, Apoe^−/−^ and DKO mice. FITC-labeled αSMA and TGFβR1 and rhodamine-labeled CD31 and DAPI (nuclei, blue) were used. Merged images are shown. Scale bar: 50 μm in each panel. Representative pictures are shown. N=5/group. Data are shown as mean ± SEM. Tukey test was used for the analysis of statistical significance. * p < 0.05.

### Inhibition of canonical Wnt signaling improves renal fibrosis

To determine whether inhibition of the Wnt signaling pathway could ameliorate the observed fibrosis, we utilized LGK974, a small molecule inhibitor of all secreted Wnts^7^. **Fig. S3a-b** depicts the schematic diagram showing the experimental protocol for LGK974 treatment in diabetic CD-1 and UUO mice. LGK974 greatly diminished the ECM deposition, relative area fibrosis, collagen accumulation and glomerulosclerosis in both models used (**Fig. S3c-d)**. Wnt inhibition significantly restored the endothelial GR level and suppressed the level of β-catenin, a marker of canonical Wnt signaling, in the diabetic and UUO mice **(Fig S3e-h)**. LGK974 significantly suppressed the elevated level of IL-1β, IL-6, IL-10, G-CSF, TNFα, MCP-1, and CCL4 whereas caused elevation in the level of CCL5 **(Fig. S4**).

### Wnt inhibitor partially suppresses the fibrogenic phenotype in the kidneys of diabetic GR^ECKO^

To determine whether Wnt inhibition could mitigate the renal fibrosis observed in diabetic mice lacking endothelial GR, a cohort of animals was treated with the Wnt inhibitor, LGK974. At the age of 8 weeks, control and GR^ECKO^ mice were injected with STZ 50 mg/kg for five consecutive days. Sixteen weeks after injection, LGK974 was administered by oral gavage for eight additional weeks (**Fig. 5a**). At the time of sacrifice, there were no differences in body weight or glucose among the groups (**Fig. 5b**). However, a significant reduction in kidney weight was observed in the Wnt-inhibitor treated diabetic GR^ECKO^ and diabetic control mice (**Fig. 5a**). Wnt inhibitor clearly improved the relative area of fibrosis, relative collagen deposition and tubular damage in the diabetic control mice; this effect was less pronounced, though still significant in the diabetic GR^ECKO^ mice **(Fig 5c**). A similar pattern was observed in the staining of fibronectin and β-catenin **(Fig 5d-e)**. Wnt inhibitor significantly suppressed EndMT (CD31/αSMA co-positive cells) in the diabetic control mice; this effect was less pronounced in the diabetic GR^ECKO^ mice **(Fig 5f)**. However, Wnt inhibition significantly reduced the level of EMT (E-cadherin/αSMA co-positive cells) in control and diabetic GR^ECKO^ mice **(Fig 5g)**.

**Figure 5.**
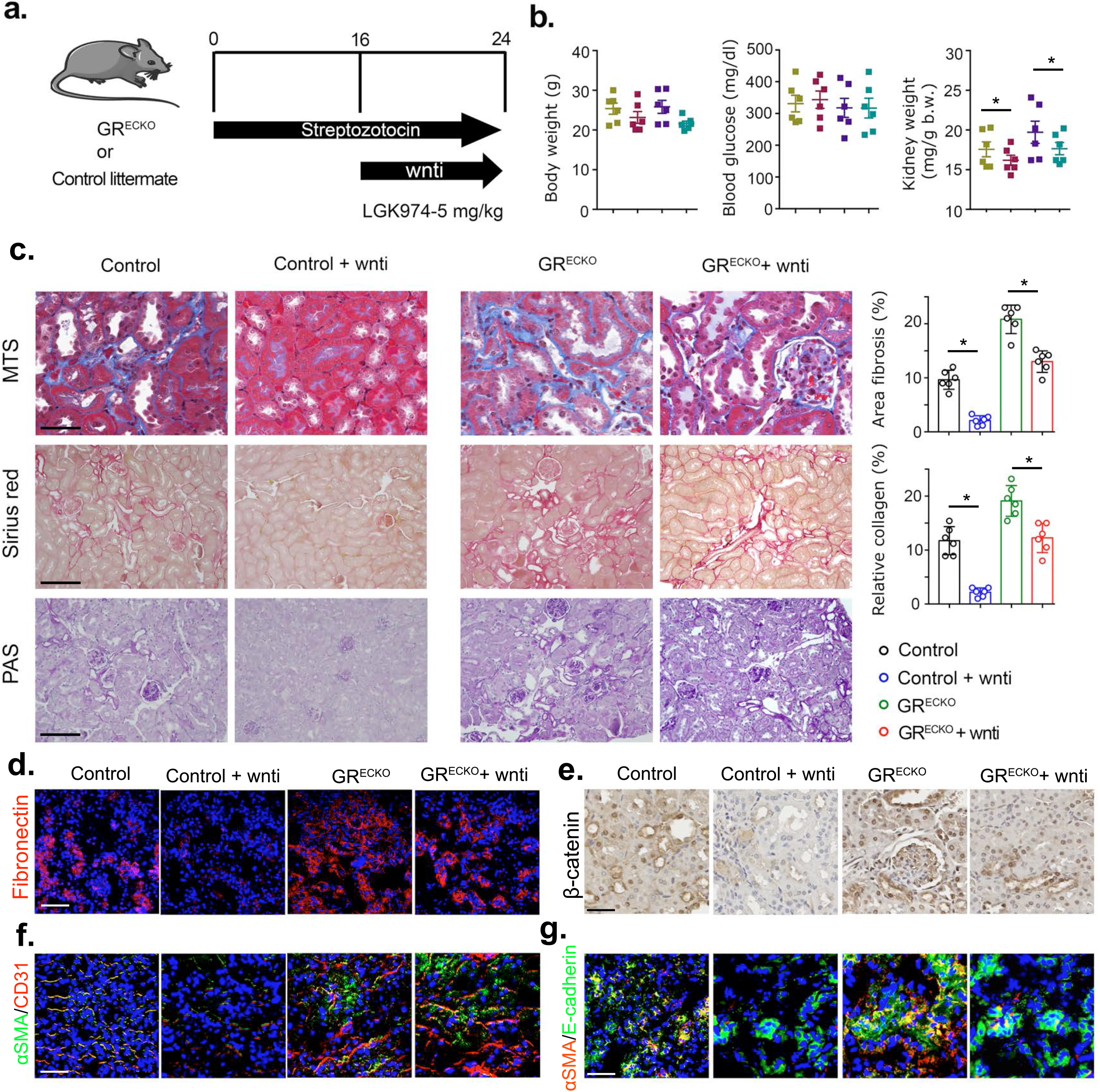
Wnt inhibitor partially abrogates the renal fibrosis in diabetic GR^ECKO^ and DKO mice. **(a)** Schematic diagram, showing the treatment protocol of wnti in the diabetic GR^ECKO^ and DKO mice. **(b)** Physiological parameters including body weight, blood glucose and kidney weight were analyzed in the wnti-treated diabetic GR^ECKO^ and DKO mice. N=6/group. **(c)** Masson trichrome, Sirius red and PAS staining in the kidneys was analyzed. Representative images are shown. Relative area fibrosis (%) and relative collagen deposition (%) were measured using the ImageJ program. N=6/group Scale bar: 50 μm in each panel. Data are shown as mean ± SEM. **(d)** Immunofluorescence analysis of fibronectin in the kidneys of wnti-treated diabetic GR^ECKO^ and DKO mice using rhodamine-labeled fibronectin and DAPI (nuclei, blue). Representative images are shown. Scale bar: 50 μm in each panel. N=6/group. **(e)** Immuno-histochemical analysis of β-catenin level in the wnti-treated diabetic GR^ECKO^ and DKO mice. **(f-g)** Immunofluorescence analysis of αSMA/CD31 and αSMA/E-cadherin were performed in the kidneys of Wnti treated diabetic control, and GR^ECKO^. In the first panel FITC-labeled αSMA, rhodamine-labeled CD31 and DAPI (nuclei, blue) were sued. In the second panel FITC-labeled E-cadherin, rhodamine-labeled αSMA and DAPI were used. Merged images are shown. Scale bar: 50 μm in each panel. N=6/group. Tukey test was used for the analysis of statistical significance. * p < 0.05.

### Metabolic reprogramming by loss of endothelial GR accelerates renal fibrosis

It is increasingly recognized that defects in central metabolism contribute to kidney fibrosis^34,41^. Defective FA metabolism in EC leads to EndMT events^42^. To investigate whether FA metabolism was deranged in our model, we performed radiolabeled [^14^C]palmitate uptake experiments in isolated EC from mouse kidneys. We observed that FA uptake was higher in isolated EC from the diabetic kidneys of the more fibrotic strain (diabetic CD-1) when compared to kidney EC from the less-fibrotic strain (diabetic C57BL/6). Administration of the Wnt inhibitor suppressed FA uptake. Kidney EC from both diabetic GR^ECKO^ and DKO mice displayed higher FA uptake when compared to that of the diabetic control littermates **(Fig. 6a)**. FA oxidation (FAO) was also assessed by measuring the ^14^CO_2_ release from radiolabeled [^14^C]palmitate in cultured EC isolated from kidneys. FAO was diminished in the isolated kidney EC of diabetic CD-1 mice and Wnt inhibitor was able to restore the level of FAO. The cultured kidney EC from diabetic GR^ECKO^ and diabetic DKO mice showed a diminished level of FAO when compared to their diabetic control littermates **(Fig. 6b)**. In the next set of experiments, diabetic CD-1 mice were treated with the FA synthase inhibitor C75, the FAO inhibitor etomoxir, the PPARα agonist fenofibrate, and the cholesterol-lowering drug simvastatin for 4 weeks. Fenofibrate and C75 ameliorated the fibrogenic phenotype, whereas etomoxir exacerbated the fibrosis. Simvastatin treatment did not cause any significant suppression in the level of fibrosis. **(Fig. S5a)**. Fenofibrate and C75 restored the level of GR protein in CD31 positive cells, whereas etomoxir and simvastatin suppressed it **(Fig. S5b)**. Fenofibrate and C75 downregulated the fibronectin and αSMA mRNA level, whereas etomoxir upregulated and simvastatin did not cause any significant change in the gene expression level of fibronectin and αSMA in the diabetic kidneys **(Fig. S5c)**. These FA modulators did not cause any significant differences in the level of blood glucose (**Fig. S5d)**. Etomoxir treatment caused significant suppression of FAO, as measured by ^14^CO_2_ release, and CPT1a level, and induced the protein expression level and β-catenin whereas, C75 and fenofibrate increased the level of FAO, induced the level of CPT1a and suppressed the level of β-catenin in the diabetic CD-1 mice (**Fig. S5e-f)**.

**Figure 6.**
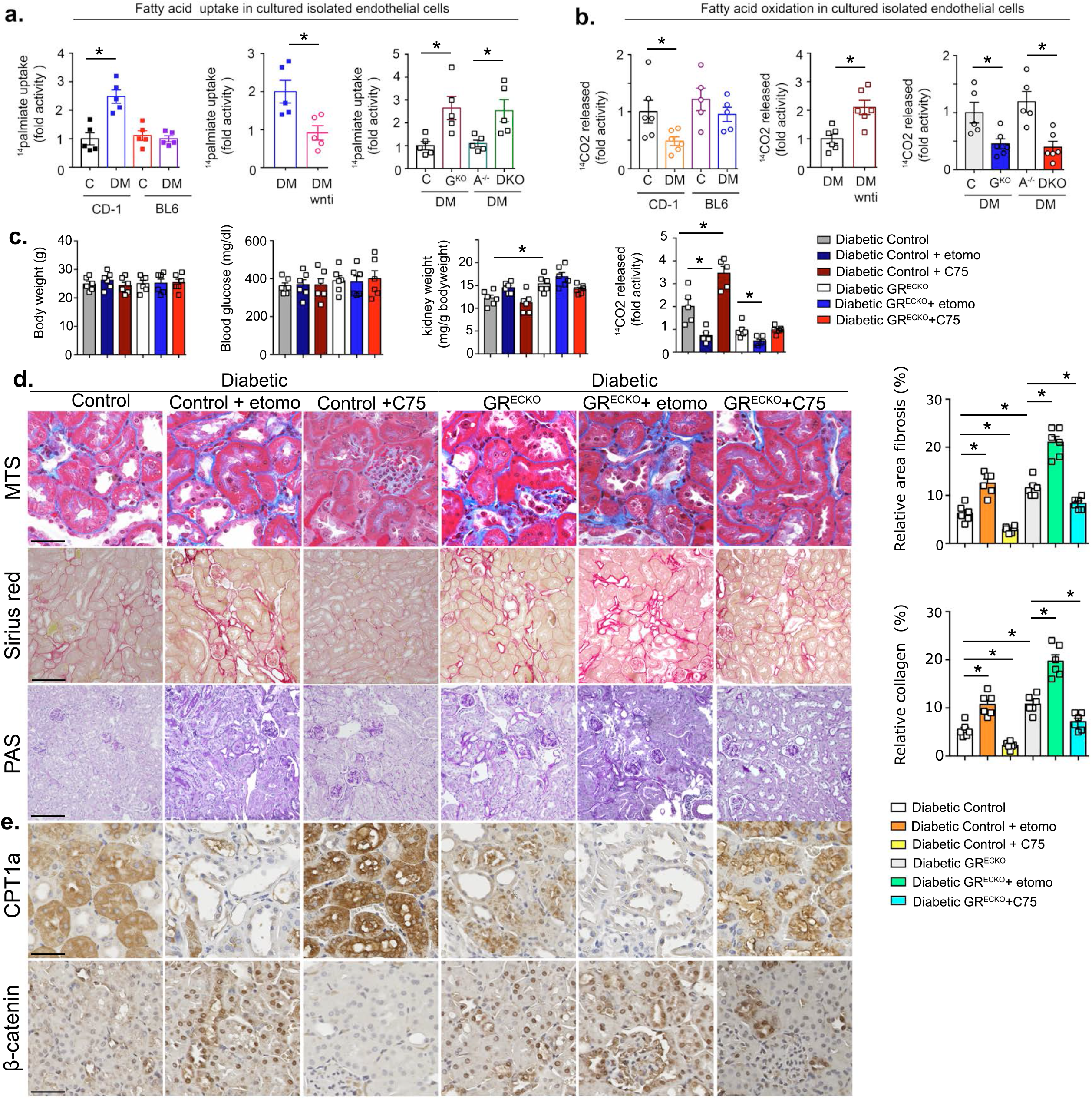
Metabolic reprogramming by loss of endothelial GR loss worsens the phenomenon of diabetic kidney disease. **(a)** Radiolabeled [1H^3^]triolein uptake analysis in kidneys in three sets of experiments 1) control and diabetic mice of CD-1 and C57BL/6 strains 2) diabetic and wnti-treated diabetic mice 3) control, GR^ECKO^, *Apoe*^*−/−*^ and DKO mice. CPM of each samples were counted. N=5/group. **(b)** Radiolabeled [^14^C]palmitate oxidation and [^14^CO_2_] release were measured. CPM of each sample was counted. N=5-6/group **(c)** Body weight, blood glucose, and kidney weight was measured at the end of these experiments. Ex vivo radiolabeled [14C]palmitate oxidation and [^14^CO_2_] released were measured. CPM of each sample was counted. N=5-6/group. **(d)** Masson trichrome, Sirius red and PAS staining were analyzed in the kidneys of control, diabetic, fenofibrate-, etomoxir-, C75-, simvastatin-treated diabetic mice. Representative images are shown. Relative area fibrosis (%) and relative collagen deposition (%) were measured using the ImageJ program. N=6/group. Scale bar: 50 mm in each panel. Data are shown as mean ± SEM. **(e)** Immunohistochemical analysis of CPT1a and β-catenin. Representative images are shown. Scale bar: 50 μm in each panel. N=6/group. Tukey test was used for the analysis of statistical significance. *p < 0.05. C-control, DM-diabetic, G^KO-^GR^ECKO^, A^−/−^-*Apoe*^−/−^

Etomoxir and C75 were also tested in the diabetic control littermates and diabetic GR^ECKO^ mice. There were no significant differences in body weight, blood glucose or kidney weight in diabetic control littermates and GR^ECKO^ mice after treatment with etomoxir or C75. Data from kidney EC revealed that etomoxir caused significant suppression in FAO, and C75 restored FAO in diabetic control littermates. However, etomoxir caused significant suppression in FAO and C75 was unable to rescue the level of FAO in kidney EC from diabetic GR^ECKO^ mice (**Fig. 6c)**. Etomoxir treatment accelerated the renal fibrogenic phenotype, suppressed the CPT1a level and increased the expression level of β-catenin in the kidneys of diabetic control and diabetic GR^ECKO^ mice. C75 treatment clearly abolished the renal fibrogenic phenotype, restored CPT1a and completely diminished the level of β-catenin in the kidneys of diabetic control. These effects were also observed in the GR^ECKO^ mice, though to a lesser extent. **(Fig. 6d)**.

### GR loss-linked EndMT disrupts central metabolism and induces mesenchymal transformation in tubular epithelial cells

This *in vivo* data suggests that the GR^ECKO^ mice exhibit enhanced EMT in their diabetic kidneys. To test whether endothelial GR deficiency affects mesenchymal programs and causes defects in central metabolism in neighboring epithelial cells, we analyzed the effects of culture media from GR knockdown HUVECs on the mesenchymal phenotype of HK-2 cells (**Fig. 7a**). Conditioned media (CM) from GR siRNA-transfected HUVECs decreased E-cadherin protein levels and increased αSMA, TGFβR1 and β-catenin protein levels in HK-2 cells when compared to media from scrambled siRNA-transfected HUVECs (**Fig. 7b-c**). CM treatment from GR siRNA-transfected HUVECs caused a significant reduction in the level of FAO, oxygen consumption rate and cellular ATP level in the HK-2 cells **(Fig. 7d-f)**. CM significantly down-regulated the level of the FAO-responsive genes *Cpt1a*, *Cpt2*, *Pparα*, and *Pgc1α* **(Fig. 7g)**.

**Figure 7.**
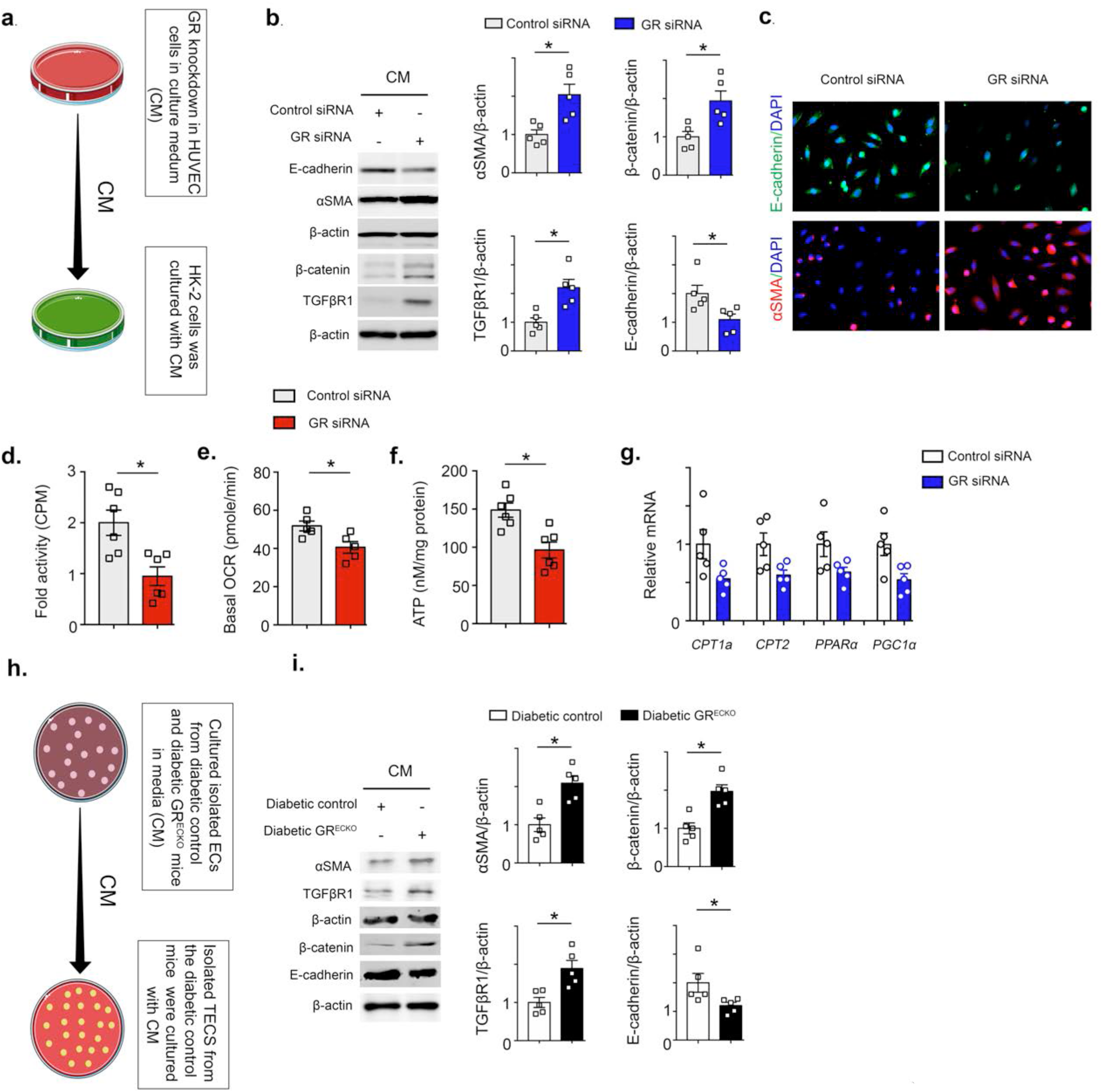
GR-loss in endothelial cells reprograms the central metabolism in the renal tubular cells, activates EMT processes. **(a)** Conditioned media experiment design. HUVECs were transfected with scrambled or GR siRNA; after 6 h, the medium was changed and cells were incubated for 72 h. The subsequently harvested media was transferred to HK-2 cells. **(b)** Representative western blotting analysis of E-cadherin, αSMA, TGFβR1 and β-catenin expression. Five independent experiments were performed. Densitometric analysis of the levels relative to β-actin is shown. **(c)** Immunofluorescence microscopy analysis of E-cadherin and α-SMA expression in conditioned medium treated TECs. For each slide, images of six different fields of view at Å~ 400 magnification were evaluated. Scale bar 30 μm. **(d)** [14C]palmitate oxidation measured by [^14^CO_2_] release. CPM were counted and normalized to the protein in the well. Three independent set of experiments were performed. **(e)** Oxygen consumption rate (OCR) in conditioned medium treated-TECs; each data-point represents the mean of eight independent samples. OCR were measured in a Seahorse XF96 analyzer. **(f)** Cellular ATP measurement. N=6 were analyzed. **(g)** Relative mRNA levels determined by qRT-PCR of regulators of FAO in conditioned media-treated TECs. **(h)** Experimental design for conditioned media. EC from the kidneys of diabetic GR^ECKO^ and diabetic control were cultured for 96 h. The subsequently harvested media was transferred to TECs from diabetic control mice. **(i)** Representative western blotting analysis of αSMA, E-cadherin, TGFβR1 and β-catenin expression. Five independent experiments were performed. Densitometric analysis of the levels relative to β-actin is shown. Data are mean ± SEM. Tukey test was used for the analysis of statistical significance. *p < 0.05.

To confirm these *in vitro* results, we isolated primary EC from diabetic control and diabetic GR^ECKO^ mice to analyze the contribution of GR-deficient EC on the mesenchymal activation in tubular epithelial cells (TECs). CM from isolated cultured EC from the kidneys of diabetic GR^ECKO^ and diabetic control littermates was transferred to cultured TECs from diabetic control mice **(Fig. 7h)**. The CM treatment from GR-deficient cells caused significant suppression of E-cadherin levels and induction of αSMA, TGFβR1 and β-catenin protein levels in TECs **(Fig. 7i)**.

## Discussion

This study demonstrates the crucial role of EC GR in the regulation of fibrogenic processes in a mouse model of diabetic kidney disease. Our results demonstrate that EC GR regulates the mesenchymal trans differentiation process by influencing FA metabolism and control over canonical Wnt signaling in the kidneys of diabetic mice. GR loss is one of the fibrotic phenotypes in diabetes that leads to disruption of cytokine and chemokine homeostasis by up regulating canonical Wnt signaling. Ultimately, these processes alter the metabolic switch in favor of defective FA metabolism and associated mesenchymal activation in TECs.

Metabolic reprogramming in EC is a crucial event in the development of myo-fibroblast formation, proliferation and fibrosis in diabetic kidneys^20,25,41,43,44^. Our data suggests GR deficiency is a critical step for the metabolic reprogramming in kidney EC. The altered cytokine levels of in the plasma of GR^ECKO^ mice include elevated levels of pro-inflammatory cytokines (IL-1β, IL-6, and IL-17) and the anti-inflammatory cytokine IL-10. The role of IL-10 has not been fully investigated in renal fibrosis in diabetic kidney disease so far. There are a few reports showing that altered cytokine levels can affect renal lipid metabolism in diabetic kidney disease^45,46^.

Recently, we demonstrated that loss of endothelial GR results in up-regulation of canonical Wnt signaling (Zhou et al, JCI Insight, In press). It is accepted that GR performs its anti-inflammatory actions by targeting the NF-kB signaling pathway^47^. However, GR targets also canonical Wnt signaling in EC which is independent of its classic target, NFkB (Zhou et al., JCI Insight, In press; ^47^). Inhibition of Wnt signaling in EC may prove to be a valuable therapeutic opportunity for combating diabetic kidney disease. The Wnt pathway is known to be an important contributor to renal fibrosis and activated canonical Wnt signaling contributes to the disruption of cytokine and chemokine homeostasis^48–51^. Our data demonstrate that higher levels of GR-deficient-associated canonical Wnt signaling are associated with the induction of mesenchymal and fibrogenic markers.

To further test the therapeutic potential of Wnt inhibition, we used the small molecule (Wnt inhibitor-LGK974). Wnt inhibition clearly suppressed canonical Wnt signaling and substantially improved fibrogenic phenotype in our mouse model of diabetic kidney disease and restored the endothelial GR level. These data suggest that GR performs its anti-fibrotic action by tonic repression of canonical Wnt signaling in EC. Notably, this effect was less evident in GR^ECKO^ possibly since Wnt inhibition was able to suppress EMT processes in other cell types (TECs) whereas, it was unable to mitigate EndMT processes. Cumulatively, these data suggest that endothelial GR is a key anti-EndMT molecule.

Research about lipid metabolism in kidney cells is limited^52,53^ but gaining importance. Defects in central metabolism contribute to diabetic kidney disease^34,41^. Clinical observations indicate a potential association between lipid levels and kidney disease^54^, and lipid control appears to be important in the prevention and treatment of diabetic kidney disease^52,53^. Here, we aimed to dissect the contribution of lipid metabolism of EC to the regulation of diabetic kidney disease. The connection between atherosclerosis and glomerulosclerosis was suggested two decades ago by Diamond, who introduced the foam cell (lipid overloaded macrophage) as the pivotal culprit in both disease processes^55,56^. The Study of Heart and Renal Protection (SHARP) study was a double-blind, placebo-controlled trial that aimed to assess the safety and efficacy of reducing LDL cholesterol in more than 9,000 patients, with or without diabetes, with chronic kidney disease^57,58^. Of approximately 6,000 patients who were not on dialysis at randomization, allocation to simvastatin therapy did not produce a significant reduction in any measure of renal disease progression^57^. There are several clinical trials on statins and their effects on kidney outcome^59–62^. These clinical data suggest that lipid lowering drugs improve cardiovascular function in diabetic patients, but do not necessarily improve the kidney outcome^53,63^. Large-scale clinical trials that are prospective, randomized, and controlled are still lacking^53^.

Based upon our previous work, DKO mice show worsened atherosclerosis, compared to *Apoe −/−* mice, which is not explained by differences in plasma lipid levels^1^. This observation was the catalyst which led to the evaluation of the diabetic phenotype in these animals. It is clear from our data that hypercholesterolemia worsened the severity of renal fibrosis in endothelial cell GR knock-out mice, suggesting that hypercholesterolemia affects EC metabolism and contributes to renal fibrosis. However, similar to available clinical data, the cholesterol lowering drug simvastatin did not ameliorate the severity of renal fibrosis in this mouse model of diabetic kidney disease. Fibrates are a class of drugs that treat hypertriglyceridemia with residual elevation of non-HDL cholesterol. However, the role of fibrates in patients with diabetic kidney disease has yet to be determined^53,64^.

We assessed the contribution of EC GR-loss linked Wnt activation and its association with defects in FA metabolism. Disruption of endothelial FA metabolism contributes to activation of EndMT in diabetic kidneys^3,4,16^,. FAO activation caused remarkable suppression of fibrosis by restoring the endothelial GR level in diabetic mice. In contrast, FAO inhibition caused acceleration in fibrosis by diminishing the level of endothelial GR in diabetic control mice, suggesting that endothelial GR is a critical protein for the action of FAO modulators. Our data clearly suggests that the antifibrotic effect of the FAO activator C75 is dependent on endothelial GR, in turn suggesting that EC are required for the anti-fibrotic action of this FAO activator.

When conditioned media from GR-deplete EC from diabetic GR^ECKO^ mice was transferred to cultured TECs from diabetic control kidneys, we observed a gain of mesenchymal markers, activation of TGFβ and canonical Wnt signaling and concomitant suppression of epithelial cell markers. These findings suggest that EndMT leads to the activation of EMT processes in diabetes. GR-deplete cells have higher levels of TGFβ-smad3 and canonical Wnt signaling, associated with disrupted levels of plasma cytokines and suppressed FAO. The cumulative effects of these metabolic changes result in activation of mesenchymal transformation in EC which appears to exert paracrine effects on neighboring TECs. The graphical figure demonstrates the functional importance of GR protein in EC homeostasis (Figure 8).

**Figure 8.**
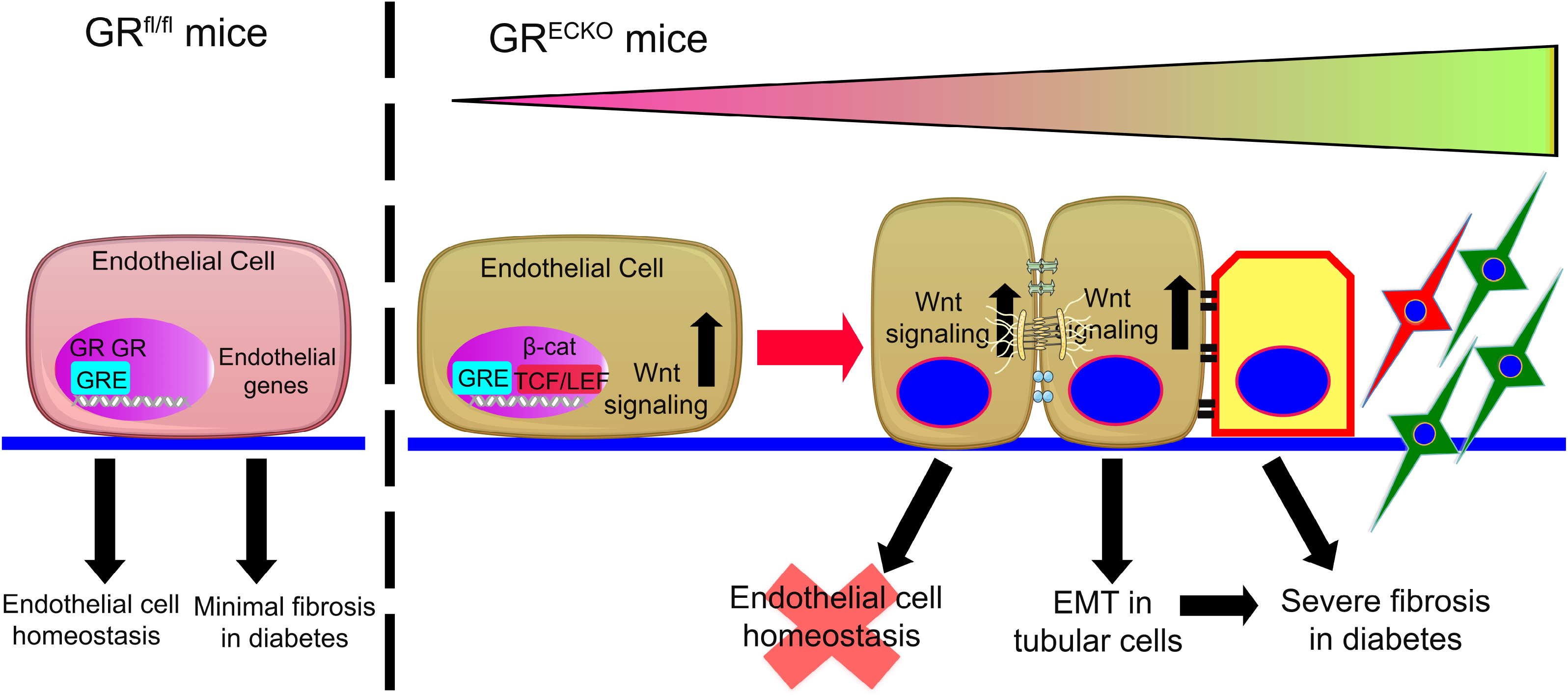
Graphical representation of the role of GR in the endothelial cell homeostasis.

In conclusion, our findings highlight the regulatory role of GR on EndMT in diabetic kidneys, mediated by control over canonical Wnt signaling and linked defective FA metabolism. This study provides new insight into the biology of GR and its critical role in renal fibrosis and diabetes.

### Limitations of study

GR agonists like dexamethasone activate GR signaling in all the cell types; however, in diabetes, dexamethasone intervention is not preferred due to the severe and predictable exacerbation of hyperglycemia. Alternative approaches which can activate GR in a cell-specific manner need to be identified and may be useful for next generation therapy for cardiovascular dysfunction and renal fibrosis in diabetic nephropathy. These data highlight the regulatory role of GR in EC. However, it is unclear whether EC GR directly regulates FAO in mitochondria or indirectly regulates this process by affecting Wnt signaling. What is the potential impact of intracellular GR translocation on the metabolic shift in EC? It will be interesting to study the role of upstream regulators of GR in different cellular compartments that might have a significant effect on disease phenotypes in the kidney. Further studies will be required to understand cellular metabolic communication in kidney disease pathogenesis.

## Acknowledgements

This work is supported by grants from the National Institutes of Health to A.D. (R01HL128406), C.F-H. (R35HL135820) and J.E.G. (R01HL131952).

## Author contributions

SPS performed experiments and wrote the paper. HZ helped in genotyping the mice and was involved in validation of the data. OS performed surgical experiments in mice. CFH and AD provided intellectual input. JG mentored, validated the data, made intellectual contributions and wrote the paper and performed final editing of the manuscript.

