## Supplemental Figures and Legends for "Loss of endothelial glucocorticoid receptor accelerates diabetic nephropathy"

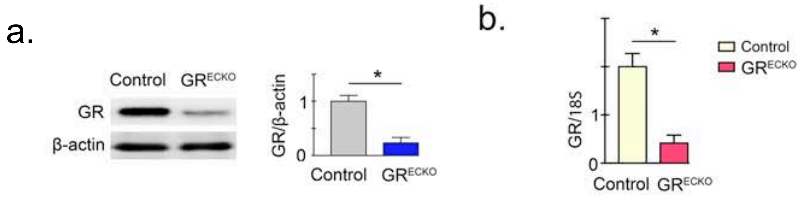


Supplementary Figure 1

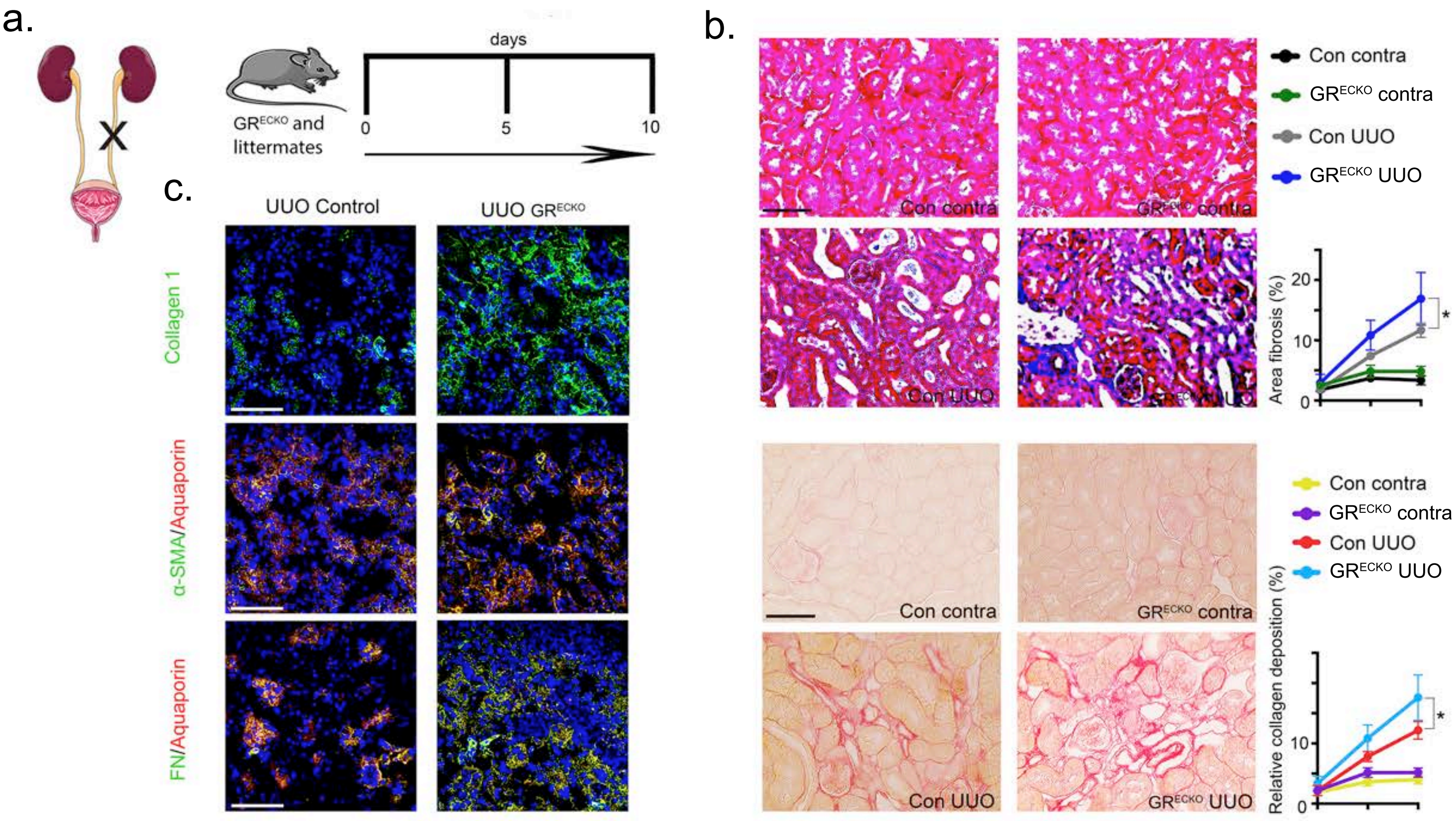


Supple Figure 2

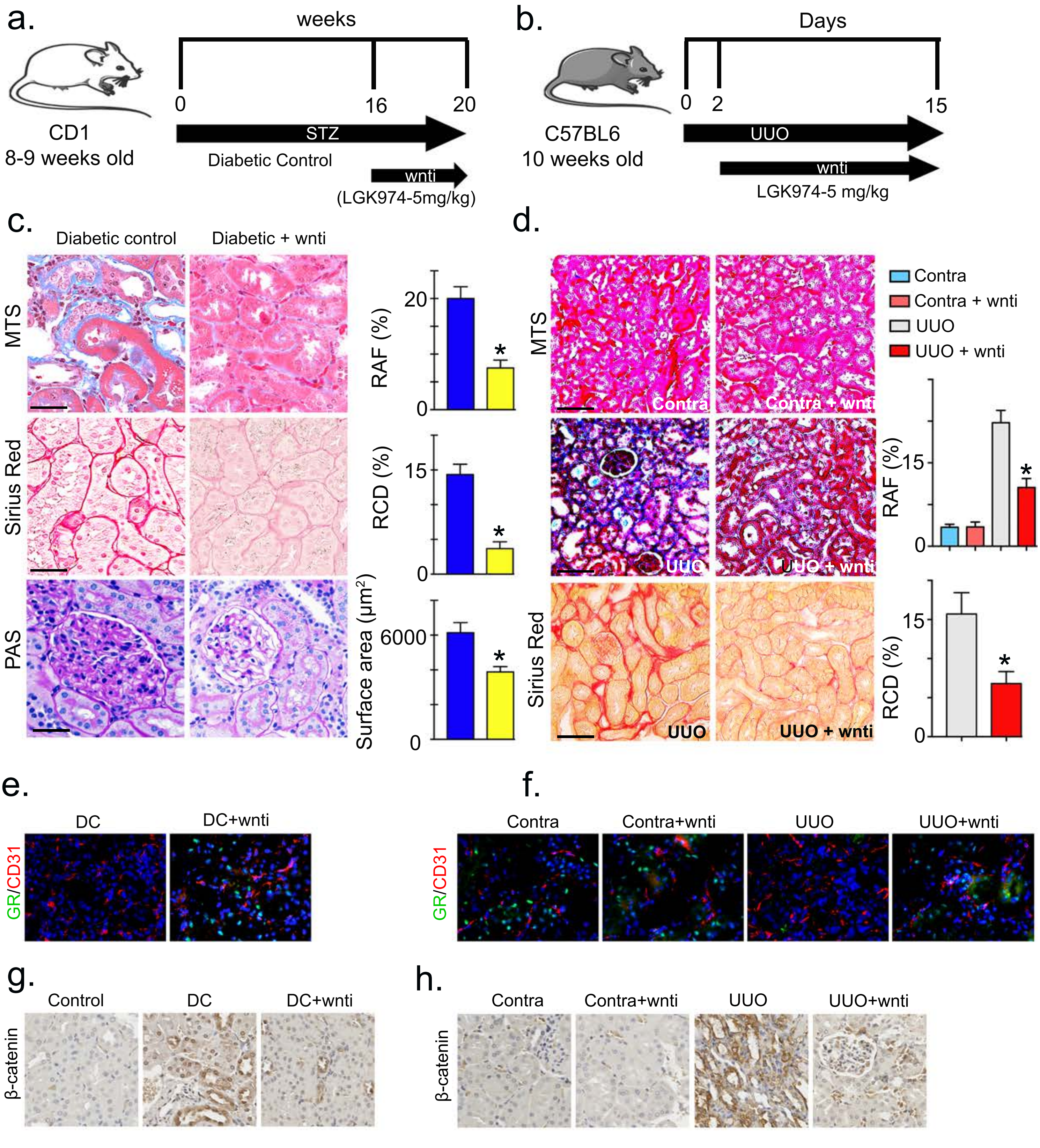


Supple Figure 3

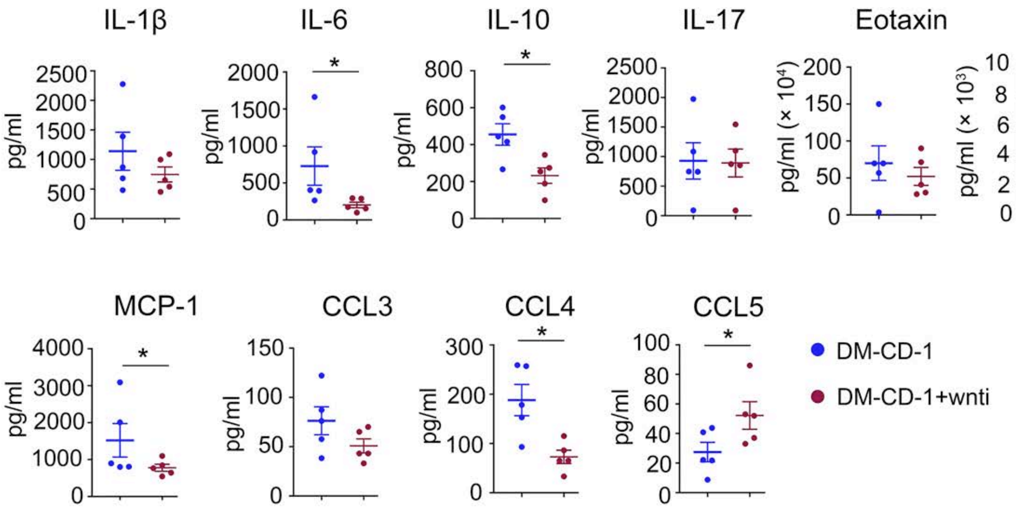


Supplementary 4

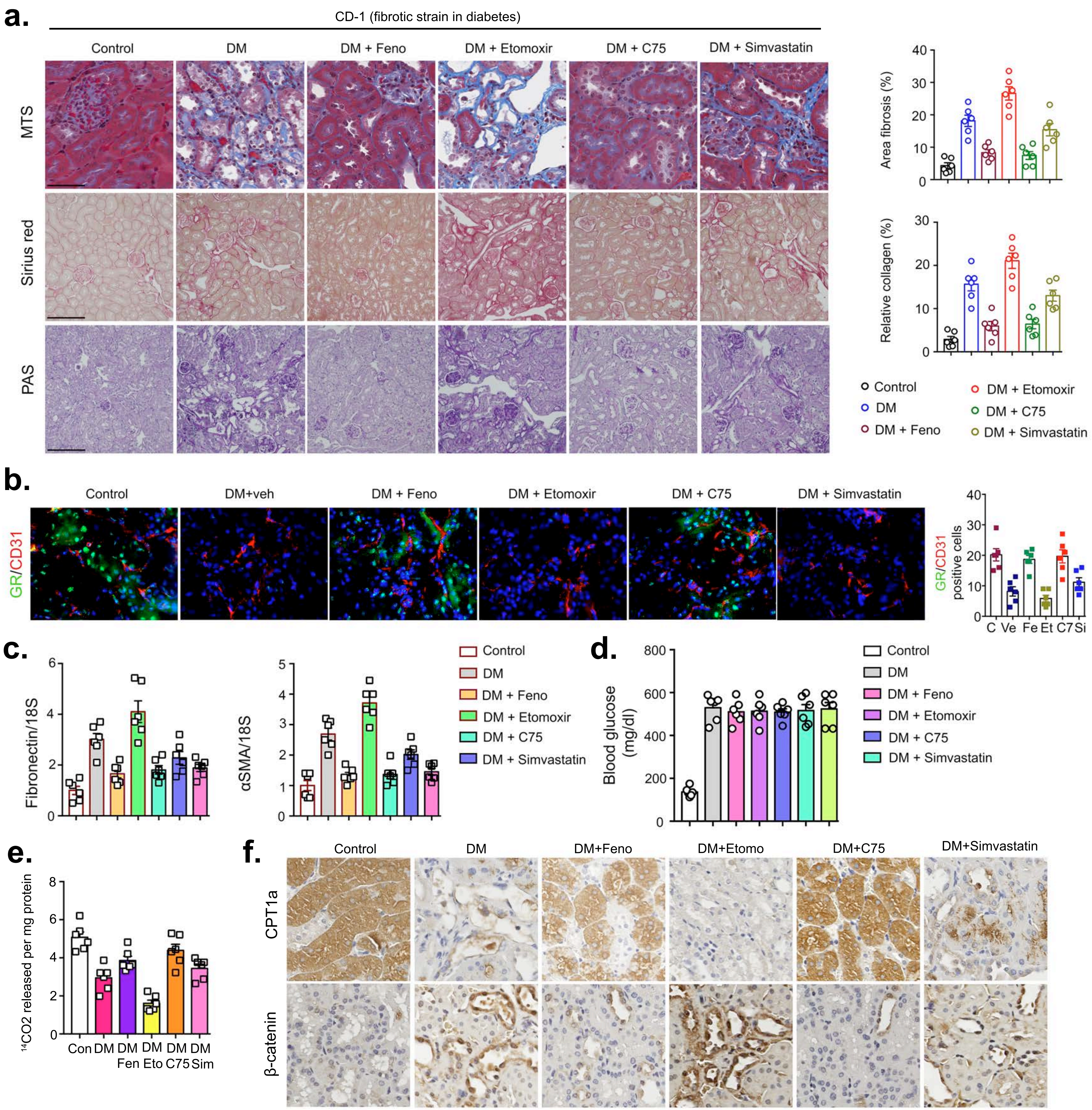

a.
CD-1 (fibrotic strain in diabetes)
Control
DM
DM + Feno
DM + Etomoxir
DM + C75
DM + Simvastatin
MTS
Sirius red
PAS
Area fibrosis (%)
Relative collagen (%)
Control
DM
DM + Feno
DM + Etomoxir
DM + C75
DM + Simvastatin
b.
Control
DM+veh
DM + Feno
DM + Etomoxir
DM + C75
DM + Simvastatin
GR/CD31
GR/CD31 positive cells
C Ve Fe Et C7 Si
c.
Fibronectin/18S
αSMA/18S
Control
DM
DM + Feno
DM + Etomoxir
DM + C75
DM + Simvastatin
d.
Blood glucose (mg/dl)
Control
DM
DM + Feno
DM + Etomoxir
DM + C75
DM + Simvastatin
e.
14CO2 released per mg protein
Con DM DM DM DM DM
Fen Eto C75 Sim
f.
Control
DM
DM+Feno
DM+Etomo
DM+C75
DM+Simvastatin
CPT1a
β-catenin

### Supplementary Figure Legends

**Figure S1. Analysis of GR protein and mRNA level in isolated EC**

**(a)** Western blot analysis of GR protein level and **(b)** qPCR analysis of GR expression level were analyzed in EC from the kidneys of control and $GR_{ECKO}$ mice. N=5/genotype. Data in the graph are shown as mean ± SEM. Student t-test was used for the analysis of statistical significance. *$p < 0.05$.

**Figure S2. Loss of EC GR worsens fibrosis in a mouse model of urinary obstruction (UUO)**

**(a)** Schematic presentation of mouse model of UUO. Left kidneys were ligated in control littermates and $GR_{ECKO}$ mice. Kidneys were excised at day 5 or 10. **(b)** Masson trichome and Sirius red staining in the contralateral and UUO-operated kidneys in control and $GR_{ECKO}$ mice was analyzed. Representative images are shown. Area fibrosis (%) and relative collagen deposition (RCD %) were measured using the ImageJ program. N=8/group. Data are shown as mean ± SEM. Scale bar: 50 mm in each panel. **(c)** Immunofluorescence analysis of collagen I, aquaporin/αSMA and aquaporin/fibronectin were performed in the contralateral and UUO-operated kidneys of control and $GR_{ECKO}$ mice. FITC labeled collagen I, αSMA and fibronectin, and rhodamine-labeled aquaporin and DAPI (blue) were used. Representative images are shown. Scale bar: 50 mm. N=7/group. Data are shown as mean ± SEM. Tukey test was used for the analysis of statistical significance. * $p < 0.05$.

**Figure S3. Inhibition of canonical Wnt signaling abolishes the fibrogenic phenotype in mice**

**(a)** Schematic diagram, representing the treatment protocol of wnt inhibitor (LGK974, at a dose of 5 mg/kg body weight) in diabetic CD-1 mice and **(b)** in UUO mice. **(c)** Masson trichrome, Sirius red and PAS staining in kidneys of diabetic and wnt inhibitor-treated diabetic CD-1 mice. Representative images are shown. Relative area fibrosis (%) and relative collagen deposition (%) were measured using the ImageJ program. N=7/group. Scale bar: 50 mm in each panel. **(d)** Masson trichrome, Sirius red and PAS staining in kidneys of UUO and wnt inhibitor-treated UUO mice. Representative images are shown. Relative area fibrosis (%) and relative collagen deposition (%) were measured using the ImageJ program. N=6/group. Scale bar: 50 mm in each panel. **(e-f)** GR protein levels in CD31-positive cells were analyzed by immunofluorescence analysis of kidneys of wnti-treated diabetic mice and wnti-treated UUO mice. FITC-labeled GR, rhodamine-labeled CD31 and DAPI blue. Merged images and representative pictures are shown. N=6/group. Scale bar: 50 mm in each panel. **(g-h)** Immunohistochemical analysis of β-catenin protein expression in wnti-treated diabetic mice and wnti-treated UUO mice. N=5/group. Data are shown as mean ± SEM. Tukey test was used for the analysis of statistical significance. *p < 0.05.

**Figure S4. Inhibition of canonical Wnt signaling disrupts the cytokine- and chemokine reprogramming in plasma of diabetic mice**

Cytokines and chemokines were measured in plasma by using the cytokine array analysis (Luminex). The plasma of wnti-treated diabetic mice was analyzed for cytokine array analysis. N=5/group. Data are shown as mean ± SEM. Tukey test was used for the analysis of statistical significance. *p < 0.05.

**Figure S5. Endothelial GR is essential for the action of anti-dyslipidemic drugs in diabetic kidney disease**

**(a)** Masson trichrome, Sirius red and PAS staining were analyzed in the kidneys of control, diabetic, and fenofibrate-, etomoxir-, C75-, and simvastatin-treated diabetic mice. Representative images are shown. Relative area fibrosis (%) and relative collagen deposition (%) were measured using the ImageJ program. N=6/group. Scale bar: 50 mm in each panel. **(b)** Co-immunolabeling of GR/CD31 was analyzed by fluorescence microscopy. FITC green-GR, rhodamine red-CD31 and DAPI (blue-nuclei) were used. Representative images are shown. Scale bar: 50 mm in each panel. N=6/group. **(c)** qPCR gene analysis of fibronectin and α-SMA in the kidneys.18S was used as the internal control. N=6/group. **(d)** Blood glucose was measured by glucometer. N=6/group. **(e)** Radiolabeled [$_{14}$C]palmitate oxidation and [$_{14}CO_2$] release were measured. CPM of each sample was counted. **(f)** Immunohistochemical analysis of

CPT1a and β-catenin in the kidneys of control, diabetic, and fenofibrate-, C75-, etomoxir-, and simvastatin-treated diabetic mice. N=5/group. Data are shown as mean ± SEM. Tukey test was used for the analysis of statistical significance. *$p < 0.05$. Con-control, DM- diabetic, feno-fenofibrate.
